# Emergent Dynamic Instability in Micrometre-scale Synthetic Active-matter Polymers

**DOI:** 10.64898/2026.07.06.736608

**Authors:** Yonatan Biniuri, Maria Bespalova, Philippe I.H. Bastiaens

**Affiliations:** Department of Systemic Cell Biology, Max Planck Institute of Molecular Physiology, Dortmund, Germany

## Abstract

Microtubules are dissipative polymers that switch stochastically between growth and rapid collapse, a behaviour known as dynamic instability. Because switching is coupled to nucleotide hydrolysis, each filament’s fate depends on the chemical state of its own subunits. Synthetic dissipative assemblies, by contrast, typically cycle as a population, following bulk fuel levels. Here we report monomer-assembled, micrometre-scale, DNA/RNA-hybrid filaments that polymerize and depolymerize autonomously over minutes. Each monomer carries an internal RNA insertion whose cleavage within the filament switches it from assembly-competent to inactive (depolymerization).

Spontaneous transesterification gives each monomer an intrinsic lifetime, so filaments grow and collapse autonomously. A DNAzyme binding-site encoded in each monomer accelerates the same RNA-cleavage up to 15-fold, raising the depolymerization rate to closely match the polymerization rate. Using this rate matching and continuous regeneration of active monomer, the system reaches a non-equilibrium steady-state in which dynamic instability emerges, with a rescue frequency of 0.7 min⁻¹ per filament. Furthermore, adding a coupling strand crosslinks the filaments into meshes whose growth accelerates autocatalytically, forming a continuously-remodelling, mesh-network. Storing the switch in each monomer lets filaments cycle independently, offering a route to autonomously remodelling active materials.

## Main

Living systems persist by continuously sensing and responding to change^1–3^. At the cellular level, this requires structures that reconfigure on timescales comparable to those of the components they organize, such as microtubules^4^ and their prokaryotic analogues^5,6^. Being rapidly-polymerizing, dissipative, and cooperative biopolymers^7,8^, microtubules remodel far-faster than the processes they direct, making them central to maintaining cellular identity and rapid adaptation^4,9^.

Microtubules assemble from α/β-tubulin heterodimers arranged in a diagonal spiral^10^. Each subunit undergoes a GTP-hydrolysis-driven conformational change^11^ that strains the polymer; when the stabilizing cap at the growing end is lost, the filament rapidly disassembles (catastrophe)^12^. Depending on the free-monomer concentration, catastrophe is often followed by a rescue event, leading to subsequent cycles of rescue and collapse^9^.

Individual microtubule filaments switch stochastically and abruptly between the two phases. These switches constitute dynamic instability, allowing them to rapidly-respond to internal and external cues. It distinguishes them from actin filaments, which predominantly treadmill, and from intermediate filaments, which are essentially stable^13–15^.

These polymer properties are essential for the many cellular processes that microtubules participate in, such as: cell polarity^16,17^, cell division^18^, cellular and organelle morpho-dynamics^8,19,20^, cargo shuttling^21^, as well as many others^10,22^.

Dynamic instability, and its essential role in living-matter identity retention^23,24^, was suggested as a key step in accomplishing the goal of a bottom-up synthetic cell formation^25–27^.

We envisaged that such a system would require: (i) growth and collapse as discrete phases that do not disrupt each other, (ii) deformable monomers that self-assemble into a supramolecular structure, (iii) an internal state within each-monomer that encodes a spatial reconfiguration and (iv) energy dissipation to sustain a non-equilibrium steady-state.

The rich toolbox of DNA nanotechnology^28–30^ offers a natural route towards such a system: nucleic-acids combine predictable-assembly properties with a broad repertoire of modifications^31^, receptors^32^, catalysts^33,34^, switches^35,36^, and templated^37,38^ and template-free^39–41^ supra-molecular structures, as well as self-assembling micrometre-long nanotubes made of repeating, monomer-like, DNA-tiles^41^, driven towards equilibrium by an enthalpically-derived lower energy state. Such nanotubes have been used to build cargo shuttles^42^, Ion-channels^43^, multiplexing systems^44^, polymer/cell hybrid structures^45^, and many more^46–48^. However, the polymer structures existed at equilibrium, and required external instructions to switch between conformational states. Similarly, chemically-fuelled reaction networks have been used to drive molecular and DNA assemblies away from equilibrium into transient, dynamic steady-states^25,26,49,50^, and dynamic-instability–like behaviour was observed in fuel-driven peptide nano-fibers^51^. Yet, in these systems, the dynamics are programmed by bulk fuel and enzyme concentrations rather than by a state stored and switched within each individual monomer. What has remained out of reach is a synthetic polymer that governs its own instability internally, growing and collapsing repeatedly, with its trajectory set by the chemical state of its subunits rather than by external inputs. Here we introduce novel meta-monomers (**MMs**) whose strands contain chemically-labile, sequence-specific RNA insertions (**Fig. 1a**), integrating intrinsic-instability into the polymer (**Fig. 1a**, inset). Cleavage proceeds spontaneously by RNA-transesterification, and shows a temperature dependence consistent with spontaneous RNA cleavage^52,53^, so destabilization requires no external switching. Furthermore, inserting DNAzyme-complementary sites in the MMs and using different DNAzyme-variants allowed for tuneable, up to 15-fold, catalytic cleavage rates.

**Figure 1:**
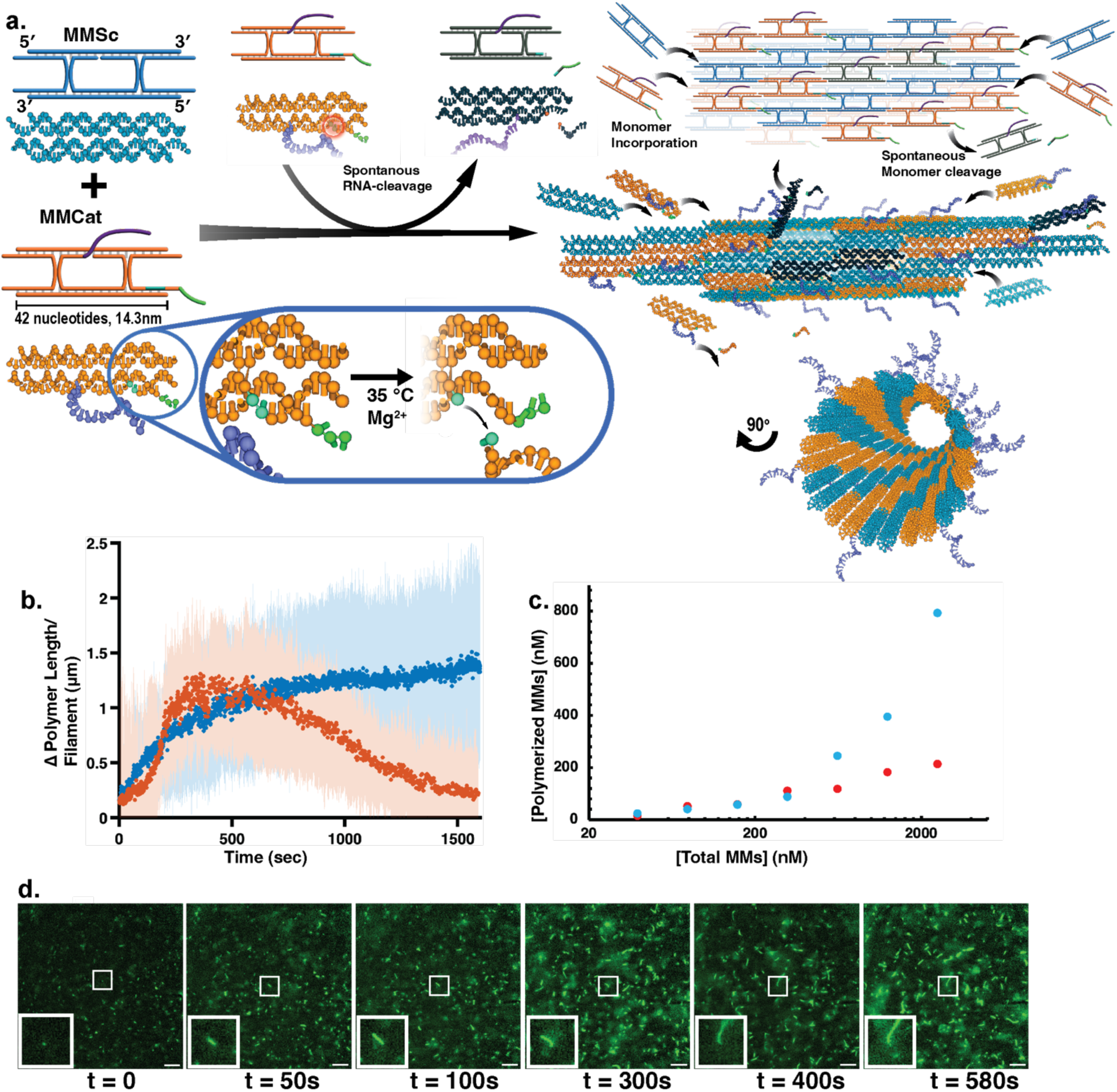
Spontaneous polymerization of DNA/RNA-monomers into μm-long polymer lattices. **a.** 2D and 3D representations of the MMSc (blue), MMCat (orange) monomers, as well as the spontaneous Mg^2+^-mediated RNA-cleavage of MMCat (dark blue) process, and self-assembly of the MMSc and MMcat into the hollow polymer filament and the and slow depolymerization mechanism originating from the spontaneously cleaving MMCat defect formation. **Below**: A 90° turned, head-on view of the hollow filament, providing a view of the externally aligned extended MMCat overhang (purple). **Inset**: Depicts the RNA inserted bases serving as the cleavage site, and the resulting spatial reconfiguration followed by RNA cleavage. Colour legend: Blue/Orange - DNA Bases, Teal - RNA bases, Green - Toehold displacement site, Purple – Outside facing overhang, Dark Blue – cleaved MMcat monomer. **b.** Time-dependent change in polymer length per filament for the all-DNA polymer (blue) and hybrid DNA/RNA MMSc/MMCat polymer (orange) at 35 ℃. Measured using Epifluorescence microscopy imaging experiments, 100 msec exposure time with 1 sec intervals between frames. **c.** Critical concentration plots for the all-DNA and DNA/RNA hybrid polymers. Monomer stocks were let incubate at 35 °C for 90 minutes. The critical concentration was observed at 35 nM for the all DNA-system, and 40 nM for the DNA/RNA system. Blue - all-DNA system, Red - DNA/RNA hybrid system. **d.** Epifluorescence microscopy time-series images of polymerizing MMSc/MMCat-mix under finite available free-monomer conditions. Stocks were mixed and immediately placed on a heated sample holder and imaged at 1 second intervals. Timelapse insets show a single filament polymerizing. scale bars: 5 μm.

The structurally encoded dissipative instability, when combined with a catalyst and an active-MMs pool, allowed for the transitioning of the filaments into a dynamically unstable steady-state.

To sustain the non-equilibrium steady-state, we used polymer-binding DNA-origami seed^46^-functionalized flow-chambers supplied-continuously with MMs, allowing for the localized, polarized growth and tracking of individual filaments over time.

Within a narrow window of competing polymerization and catalytic depolymerization rates, filaments become dynamically unstable, switching repeatedly between growth and collapse. Furthermore, when a coupling strand was added to the system, polymers clustered and captured one another, forming dynamic, autocatalytically-accelerating filament meshes.

### Hybrid DNA/RNA polymers are capable of controllable polymer growth and spontaneous RNA-cleavage-dependent collapse

Structurally encoded, dissipative instability was inserted in the monomer/polymer structure using novel, hybrid MMs containing sequence-specific RNA base-insertions. RNA forms a cyclic compound between its ribose C_2_-OH and a downstream phosphate, leading to irreversible inter-nucleotide bond-cleavage^54^. In Mg^2+^-rich environments, the Mg^2+^-stabilized C_2_-Phosphate ring-formation results in fast, spontaneous, RNA-cleavage^52^. To implement the dynamic synthetic polymer, MMs based on the DAE-E tile^55,56^ were designed. These MMs, composed of five nucleic-acid-strands, can be assembled in a wide range of spatial configurations^41^. Each monomer unit contained four complementary 5bp-long sticky-ends, programmed to form a diagonal-clockwise polymer-lattice configuration (Figure S1a). Banded and anti-clockwise configurations were also examined.

Different MM variants were used: (i) **MMSc**, acting as scaffold and fluorescent-marker and (ii) **MMCat**, acting as an intrinsic-filament destabilizer (**Figure 1a**). Using two different MM units to compose the polymer doubled the number of outer-facing nucleic-acid strand-ends, and allowed for signal multiplexing. The spontaneously-cleaving, polymer-destabilizing RNA-bases were introduced into MMCat, adjacent to one of its 5-bp duplex forming sticky-ends (**Figure 1a, inset**). This allowed for spontaneous polymer-destabilization by the irreversible removal of one of the duplex-forming sticky-ends within the MMCat unit downstream from the cleavage site. It was assumed that cleavage events, or spatial defects, that accumulated within the nucleic-acid polymer-lattice lead to a meta-stable state and the eventual collapse of the polymer.

The MMCat/MMSc DNA/RNA polymer construct used throughout this work spontaneously polymerized at a rate of 0.26 ± 0.80 µm/min using 1xTAE-Mg^2+^ buffer, 0.8 µM MMSc/MMCat in a 1:1 ratio at 35 °C (**Figure 1b**), with a persistence length of 9.43 ± 1.83 µm (Figure S1b). The rate of polymerization depended on the free initial monomer concentration (Figure S1c), with critical concentrations determined as 40 nM monomers for the DNA/RNA hybrid polymer and 35 nM for the all-DNA system (**Figure 1c** and **Figure S1d**). These conditions also allowed for proofreading within the polymer, effectively melting the mismatched, less-stable monomer duplexes. When MMCat/MMSc polymerization was attempted at temperatures of 23 °C and below, disordered aggregates formed from the DNA/RNA MMCat/MMSc-mixture, with no ordered polymerization observed (**Figure S2a**). The all-DNA system showed very slow (hours) polymerization rates, with no aggregates forming (Figure S2b). Filament polymerization (and depolymerization) was then tracked over time, **Figure 1d**. The polymerization process consisted of a slow monomer-addition and a faster oligomer incorporation step, joining at the polymer edges, similar to microtubules^57^ (**Figure 1d, insets**). After ca. 10 mins the polymer reached equilibrium (**Figure 1b, 1d**), with the DNA/RNA hybrid polymers slowly depolymerizing over the next few hours (Figure S3). The all-DNA composed polymers remained stable for time periods of up to 90 days (Figure S4). It should be noted that the filaments exhibited a lognormal distribution (Figure S5).

The hybrid DNA/RNA polymers displayed RNA cleavage-mediated degradation of the polymer structure, and the subsequent fracturing and disassembly of the micrometre-long filaments. The internally-derived depolymerization rate was measured to be 0.09 ± 0.03 µm/min at 35 °C under depleted free-monomer conditions, to ensure the absence of free-monomer available that would compete with the depolymerization reaction.

To assess the relation between filament depolymerization and RNA-cleavage, the cleavable-MMCat sticky-end was fluorescently-labelled using Atto647, and the decrease in Atto647 intensity across the filament was measured under depolymerizing conditions (Figure S9). Indeed, a decrease in the Atto647-cleavage-marker intensity was detected prior to filament collapse, demonstrating a direct link between RNA-cleavage and depolymerization.

The newly developed MMSc and MMCat units enabled the successful crossing-over of the micrometre-scale, active-matter polymer system into a dissipative state, requiring a constant energy source, uncleaved MMs, to sustain the polymerized state.

### Targeted lattice destabilization leads to controlled polymer collapse with tuneable properties

To precisely tune the RNA cleavage-kinetics rates beyond the native-instability of the hybrid-polymer, amplification of the RNA-cleavage properties using DNAzyme/polymer coupling was assessed (**Figure 2a**). The RNA-cleaving, Mg^2+^-dependent DNAzyme, **Dz**, utilized two programable filament-binding domains, and a short catalytic loop^58^.

**Figure 2:**
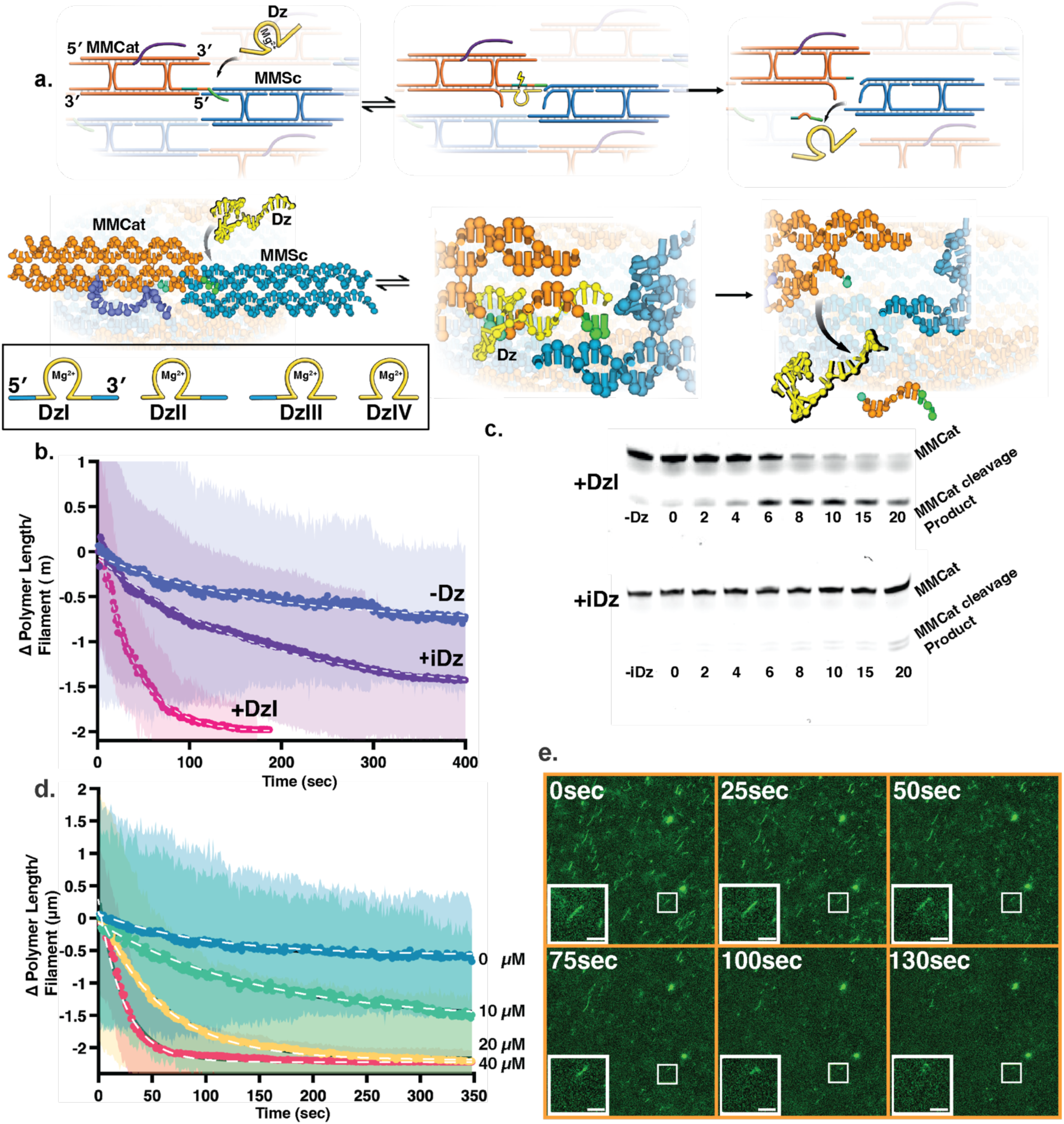
DNAzyme catalysed cleavage of DNA/RNA hybrid polymer filaments. **a.** 2D and 3D representations of the DNAzyme catalyst, DzI, invading into the polymer using the toehold-flanked active site inserted on MMCat, and subsequent catalyst/active site interaction and resulting catalysis-derived RNA-cleavage of the MMCat-unit, and removal of the MMSc-duplexing arm, leading to filament destabilization and depolymerization.(orange: MMCat, teal: MMCat RNA-insertion, green: MMCat catalyst-compatible toehold strand, purple: outside facing overhang, blue: MMSc, yellow: DNAzyme catalyst(DzI-DzIV)).**Inset**: Different DNAzyme catalyst variants used in this work: DzI, complete binding site, DzII, 5′-truncated binding site, DzIII, 3′-truncated binding site and DzIV, doubly 5′ and 3′-truncated binding sites. (Yellow: catalytic core and essential binding domain regions, Blue: Tuneable binding domain regions) **b.** Change in polymer length per filament as a function of time under monomer-depletion conditions at 35 °C in the presence of: (i) no DNAzyme, (ii) the mixed catalytic core DNAzyme variant iDz ,10 μM, and (iii) the compatible DNAzyme variant DzI, 10 μM. **c.** Top: Denaturing Urea-PAGE of the catalysed, RNA-cleavage derived polymer-degradation in the presence of saturating concentrations of the DzI catalyst over 20’ at 35 ℃. Upper band corresponds to the complete RNA-containing substrate, MMCat, and the lower band is the transesterification product. Bottom: Denaturing PAGE of the polymer in the presence of iDz, the DNAzyme lacking binding sites compatible with the MMCat active site. **d.** Change in polymer length per filament as a function of time under monomer-depletion conditions at 35 °C in the presence of 0, 10, 20, and 40 μM of the DzII catalyst. **e.** Epifluorescence microscopy time-series images depicting the DzI catalysed polymer degradation under monomer depletion conditions, scale bars: 5 μm

Different approaches to attenuate the Dz-derived catalytic-depolymerization rates were examined: (i) Varying lengths and sequence-compositions of the binding domains flanking the catalytic core in Dz, yielding varying K_M_ and *k_cat_* values, and resulting in four Dz variants: DzI, containing the complete binding domain, DzII containing a 5′-truncated binding domain variant, DzIII containing a 3′-truncated binding domain variant, and DzIV, a doubly 5′+3′-truncated variant (**Figure 2a, inset**; Figure S6). (ii) Different length and sequence compositions of the RNA cleavage-site, leading to catalyst/substrate interactions with varying kinetic properties (Table S7). (iii) The incorporation of toehold-sequences of different lengths within the MMCat cleavage site, assuming different lengths of toehold-mediated duplexes forming on MMCat/MMSc would yield catalytic reactions with varying efficacies (Table S8).

Using these different Dz(I-IV) variants, toeholds, and RNA-insertions, different catalytic rates were measured, and a range of ca. 15-fold catalytic activity was established.

To assess the catalytic function of Dz(I-IV) on the DNA/RNA hybrid-polymers, the MMCat and MMSc monomers were mixed and annealed overnight in 1x TAE-Mg^2+^ buffer at a concentration of 5 µM, resulting in a polymer-filament solution.

DzI was added at a saturating concentration (10 µM) to the annealed-polymers (25 nM), and the polymer/DzI-mixture was transferred onto a passivated glass-slide sandwich, and imaged using a heated-epifluorescence microscope at 35 ℃. The polymers depolymerized rapidly from their ends, (**Figure 2b**, curve +DzI, Movie 1) when compared with the system without catalyst (**Figure 2b**, curve -DzI, Movie 2).

Similarly, when the polymer solution was mixed with a DNA sequence containing a randomly mixed catalytic-core and complementary binding-sites, Mut, (Figure S10), or a DNAzyme with mismatching binding-sites, iDz, (**Figure 2b**, curve +iDz, Movie 3), minimal cleavage-activity enhancement was observed. This was attributed to the destabilization of the polymer by the successful-polymer-invasion of Mut, or to unspecific electrostatic interactions with iDz. The catalytic-depolymerization rate measured for DzI was *V_max_* = 114.2 ± 10.2 nM/min at 35 ℃.

The Dz(I-IV)-catalysed depolymerization-reaction was further corroborated using Urea-PAGE, and observing the evolution of the cleaved-off MMCat units over time (**Figure 2c**). We then attempted mixing 0.4 µM of separately-annealed MMSc and MMCat stocks with varying DzI concentrations. The spontaneously polymerizing monomers, when mixed with 0.3-10 µM DzI catalyst, led to the formation of very short (ca. 0.5 µm) transient structures which depolymerized entirely in the order of seconds. These results indicated the necessity for a catalyst with kinetics matching those of the MMCat/MMSc-mix polymerization rate. Kinetic analysis performed on the DzII, DzIII, and DzIV catalyst constructs showed reduced catalytic activity across all variants, with the rates of RNA-cleavage measured being *V_max_*(DzII) = 45.1 ± 6.2 nM/min, *V_max_*(DzIII) = 21.9 ± 4.1 nM/min, and *V_max_*(DzIV) = 17.8 ± 4.3 nM/min. The measured catalytic rates suggested that DzII possessed kinetic properties, compatible with the polymerization rate measured for the DNA/RNA hybrid polymer (**Figure 2d**). Mixing of 0.4 µM MMCat/MMSc with 1.2 µM DzII resulted in the activation of growth and collapse on individual polymers. Nevertheless, the reaction was tightly regulated by the amount of active monomer, resulting in a single recovery event or none, until complete depletion of the MMs (Figure S11). This property of energy consumption allowing for the spatio-temporal state-transition of monomers within the polymer, is reminiscent of GTP hydrolysis along the length of polymerizing microtubule filaments, and their nonlinear dynamics.

### Dynamic instability emergence in hybrid DNA/RNA active matter under a constant monomer flow

As polymerization is dependent on available MMCat/MMSc concentration, and breaks rapidly with MMCat consumption, we sought to examine the dynamics observed under the non-equilibrium conditions over extended time-scales. We assumed that since the dynamics are closely coupled with monomer-concentration and the Dz-catalyst activity, constant regeneration of the active-monomer available would form a non-equilibrium steady-state for longer time periods (**Figure 3a**).

**Figure 3:**
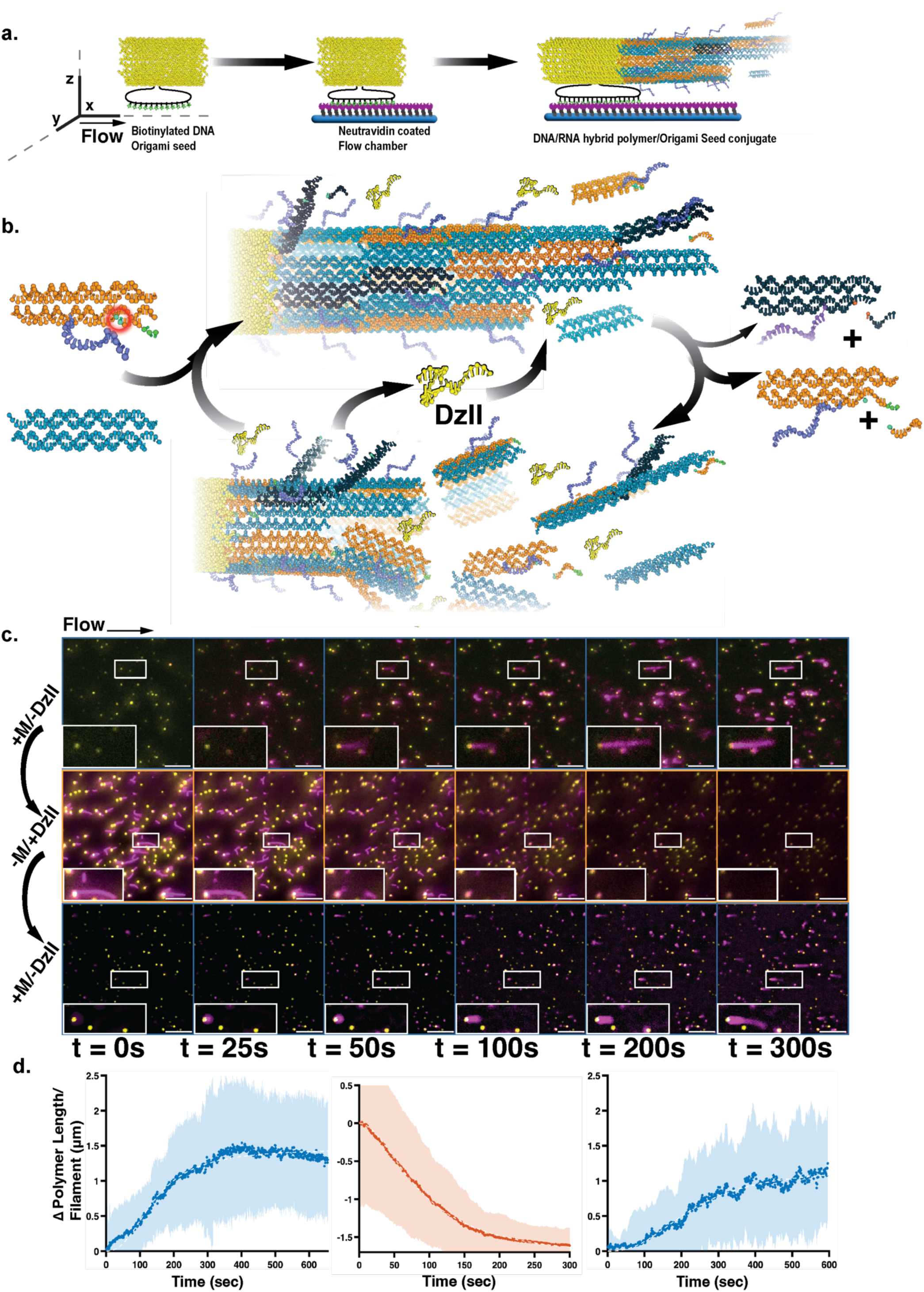
State transition in polymers between growth, collapse and dynamic instability using constant RNA-cleaving active MM-mix/DzII flux. **a.** Schematic representation of the flow chamber glass-surface coating using DNA-origami seeds, capable of binding and individual filament growth. The glass was passivated and coated using the step wise additions of biotinylated-PEG followed by Neutravidin, resulting in immobilized biotinylated-seeds on the glass surface. **b.** 3D representation of the constant consumption and supply of the MMCat and MMSc units under constant MMCat/MMSc and DzII flux in the flow chamber, allowing for the extended observation of the dynamically unstable phase of the system. **c.** Time-series images depicting the different states of the MMCat/MMSc mix as a function of different monomer and DNAzyme compositions. Top:(*+M/-DzII*) 5 ± 1 μl/min monomer solution only, leading to rapid filament population growth. Middle:(*-M/+DzII*) 5 ± 0.5 μl/min of the DzII-catalyst solution only, resulting in rapid filament collapse. Bottom:(*+M/-DzII*) 5 ± 1 μl/min monomer solution only, following a DzII induced filament population collapse. The re-introduction of active-MMCat/MMSc mix lead to rapid filament population growth. **d.** Change in polymer length per filament as a function of time under varying flow chamber conditions: Right, 5 ± 1 μl/min monomer solution only (*+M/-DzII*), Middle: 5 ± 0.5 μl/min of the DzII-catalyst solution only (*-M/+DzII*), Left: 5 ± 1 μl/min monomer solution only (*+M/-DzII*), following a DzII induced filament population collapse. scale bars: 5 μm.

To achieve a constant MMCat/MMSc-mixture concentration in solution, a multi-channel-fed flow chamber, consisting of a mixing-inlet fed from two individually controlled channels, was employed. The field of view in the chamber was passivated, coated, and functionalized with DNA-origami seeds capable of localized duplex formation with the MMs and polymer units. This allowed for the polarization and localization of individual filaments, and enabled the dynamic properties of individual filaments to be tracked and quantified (**Figure 3b**).

The flow chamber experiments were done using a custom built TIRF microscope. The shallow, 200-300 nm deep evanescence-field allowed for the use of high MMCat/MMSc concentrations and imaging at second-long intervals for ca. 200 minutes. Upon introducing a stock solution of 0.8 µM [MMCat/MMSc-mix] into the Origami-seed-labelled chamber at 5 ± 1 µl/min, seed-polymer conjugation events were observed. The polymerization rate measured was 0.3 ± 0.1 µm/min (**Figure 3c**, top time-series, **Figure 3d**, right panel, Movie 5).

At fast flow regimes of 6 µl/min to 20 µl/min, significant turbulence and shear forces were introduced along the seed-polymer conjugates, evidenced by the formation of turbulence “cones” along individual polymers^45^, leading to reduced polymerization rates, loss of signal as polymers washed-away from the evanescence field, and skewed kinetics. Transitioning to lower flow rates, 5 µl/min to 1 µl/min, allowed for more-consistent dwelling of individual polymers on the glass surface.

After ca. 10 minutes, the DNA/RNA hybrid-polymers reached a steady-state population size and length distribution, and maintained these ratios for as long as the active-monomer mix was flowed into the chamber. When the flow was substituted for 1x-TAE-Mg^2+^ buffer solution, the polymers depolymerized slowly and reduced in size and quantity over 20 minutes (Figure S12). Replacing the solution again to the active-monomer mix solution led to the rapid recovery of the polymer population, Figure S13. In contrast, the all-DNA polymer-seed conjugates exhibited higher filament stability under flow and similar conditions of the absence of the All-DNA-monomer-mix (Figure S14). To further examine the nonlinear dynamics of the hybrid DNA/RNA polymer under steady state conditions, the buffer channel was substituted to a solution containing 1.2 µM [DzII]-catalyst and the MMCat/MMSc-mix channel was stopped (**Figure 3c**, middle time-series, **Figure 3d**, middle panel, Movie 6). The total flow of the chamber was kept at 5 µl/min. Upon addition of the DzII-catalyst to the steady-state polymer-filament population, rapid collapse of the filaments was observed, indicative of catalytic RNA-cleaving activity across the polymer filaments observed.

Total, DzII-driven, filament depolymerization from their seed-anchors was observed in ca. 200 seconds, with a depolymerization rate of 0.5 ± 0.2 µm/min. To assess the recoverability of the system, the DzII channel was replaced with the MMCat/MMSc-mix once more. Indeed, individual filaments conjugated and regenerated from the Origami-seeds, yielding a new steady state filament population (**Figure 3c**, bottom time-series, **Figure 3d**, middle panel, Movie 7).

When the polymer population was mixed with the incompatible-DNAzyme, iDz, or the inactive Dz-sequence, Mut, polymer growth dominated (Figure S15).

Having established the system’s ability to maintain the growth and collapse phases separately, we sought to have both phases exist simultaneously. Solutions of 0.8 µM [MMCat/MMCat-mix] and 1.2 µM [DzII] were loaded onto separate channels, and introduced in concert into the DNA-origami-seed functionalized flow chamber. Upon mixing of the two solutions in the flow chamber, rapid growth and depolymerization cycles were observed. While most of the filaments went through a single growth and collapse cycle, 10% of the filament-population sustained multiple cycles of growth and collapse, or dynamic instability-like behaviour (**Figure 4a, 4b,** Movie 8). After ca. 400 seconds, the system achieved a new steady state, which consisted of shorter filaments exhibiting growth and collapse cycles (**Figure 4c**, **Figure 4d**). Stopping the flow of active MMs mix led to the rapid collapse of the filament population, while inversely, stopping the flow of the DzII catalyst led to the increase in filament population length, and reduced depolymerization events, demonstrating the dependence of the non-equilibrium steady-state on MMCat/MMSc and DzII.

**Figure 4.**
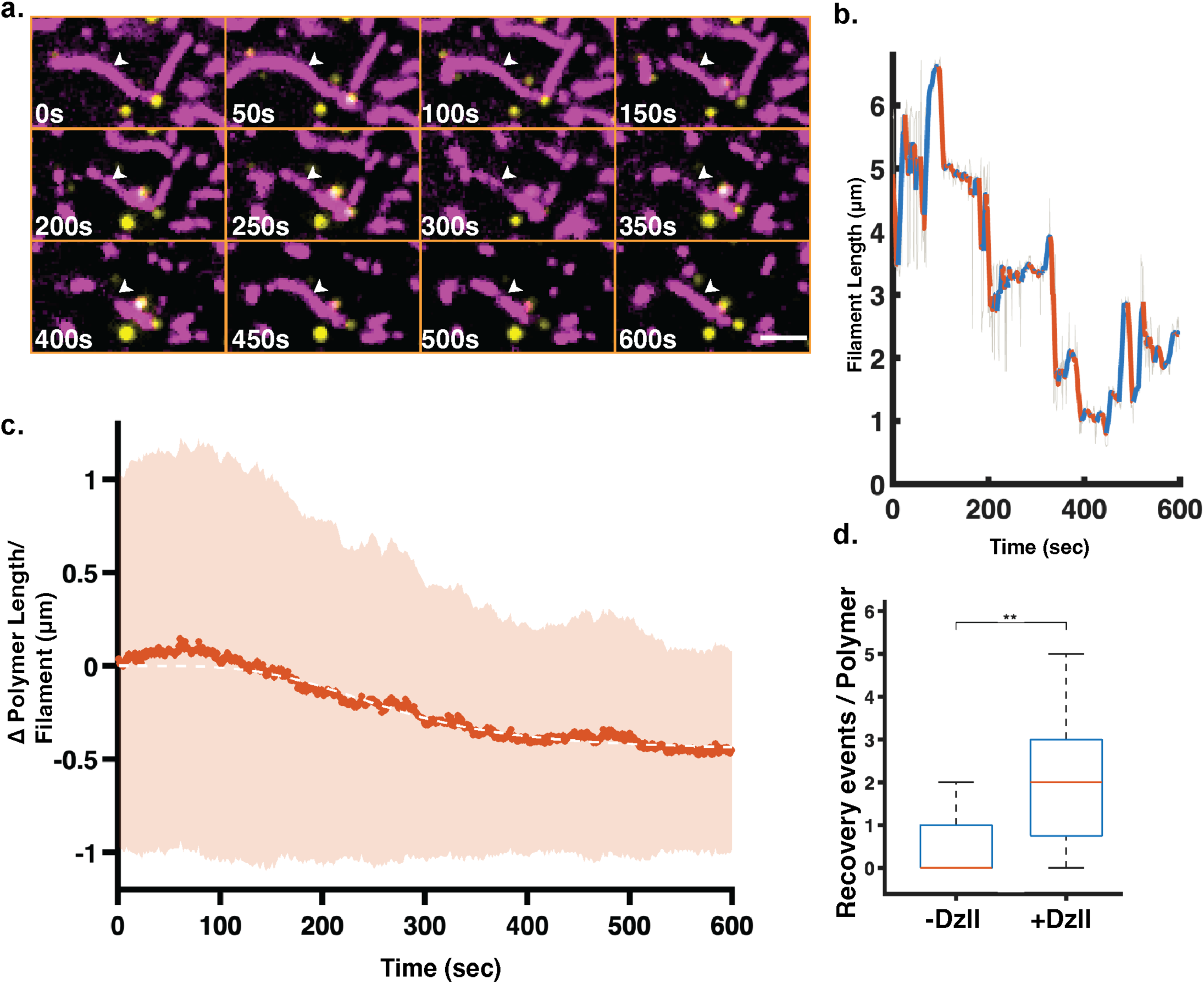
Polymer-coupling interactions lead to an auto-catalytic mesh formation capable of reinforcing a bulk identity. **a.** Time-series images of the MMCat/MMSc mix, 2.5 ± 0.5 μl/min, and the DzII, polymer compatible, RNA-cleaving catalyst, 2.5 ± 0.5 μl/min, mixed phase within the flow chamber. The mixture of MMCat/MMSc and DzII led to the formation of a dynamically unstable phase, resulting in rapidly polymerizing and depolymerizing filaments. White arrow corresponds to the filament depicted in panel **b.** Purple: filament, yellow: DNA-origami seed. **b.** Change in polymer length for a single filament as a function of time within the flow chamber for the mixed MMs/DzII conditions. Blue: filament growth, red: filament collapse. **c.** Change in polymer length per filament as a function of time within the flow chamber for the mixed MMs/DzII conditions. **d.** Box plots quantifying the change in recovery event probabilities with and without DzII in the flux of the flow chamber N = 309 for -DzII, and N = 112 for +DzII. Scale bars: 2 μm.

### Polymer-coupling interactions lead to an autocatalytically-forming dynamic mesh networks

We further investigated the crowding and coupling of individual polymer filaments into active meshes. A coupling inducing strand, **Cpl**, capable of crosslinking monomer/polymer units using the MMCat outward-facing 3′-end strand, was designed (**Figure 5a**).

**Figure 5.**
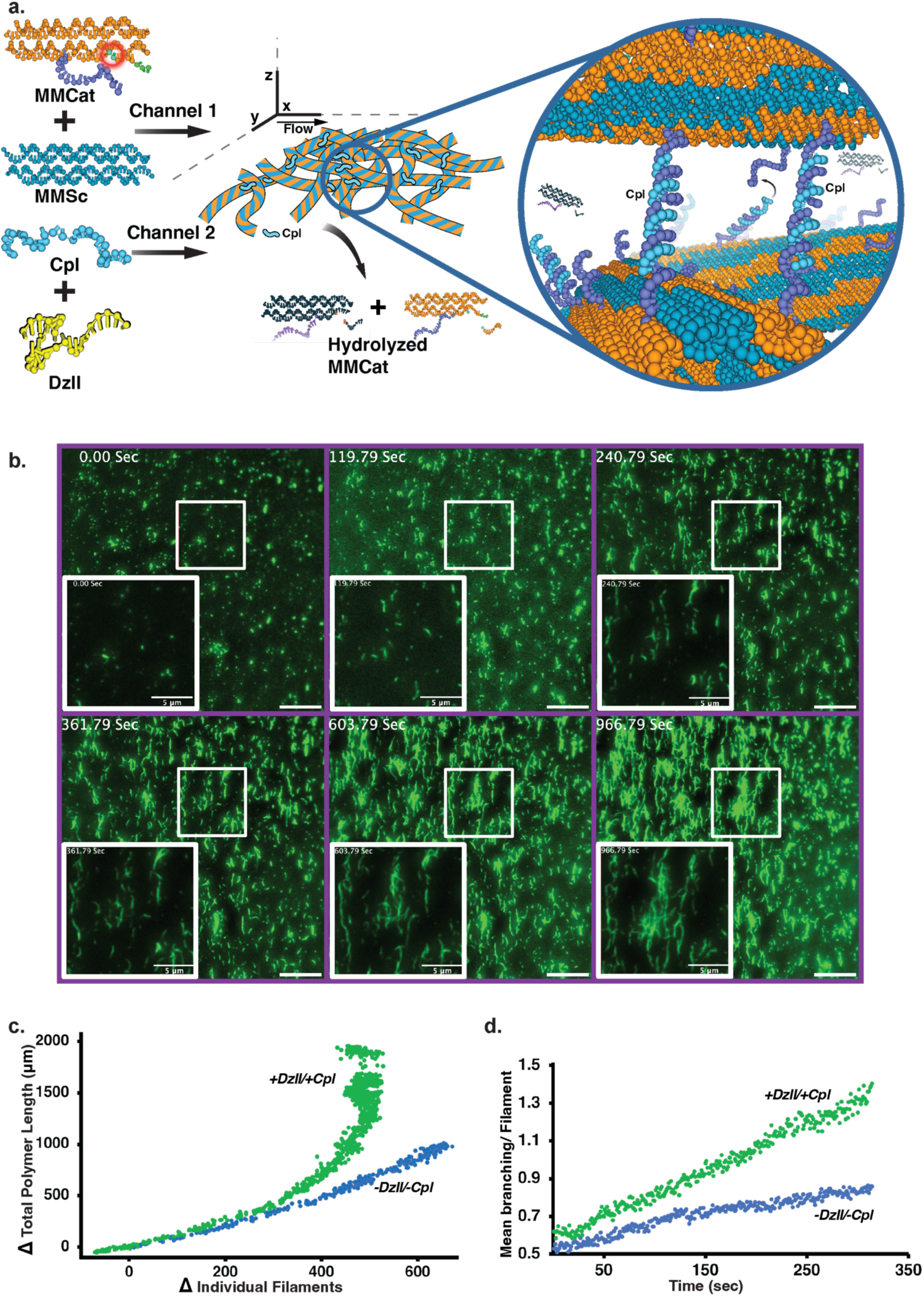
Polymer-coupling interactions lead to an auto-catalytic mesh formation capable of reinforcing a bulk identity. **a.** Schematic representation of the mixtures used in the flow chamber channels, and the resulting autocatalytic polymer-mesh. Channel 1 was loaded with MMCat/MMSc mixture ,0.8 μM, and channel 2 was loaded with the Catalyst DzII, 1.2 μM, and the coupling strand Cpl, 4.2 μM. Inset: Detailed representation of the DzII/Cpl-induced polymer-meshing interaction, by their direct duplex formation **b.** Image series depicting the evolution of the mesh-forming, active-matter polymers over time. **c.** Total polymer length as a function of individual seeded filaments within the flow chamber for DzII/+Cpl (green), and -DzII/-Cpl (blue) system configurations. **d.** Time dependent mean-change in the branching per filament for DzII/+Cpl (green), and -DzII/-Cpl (blue) system configurations.

It has been previously shown that microtubules filaments exhibit properties of self-capture and clustering^8^. We envisaged that similarly, the synthetic active-matter MMCat/MMSc polymer could form meshes capable of a network-function (**Figure 5a, inset**). By complementing the MMCat/MMSc/DzII-mix with Cpl, individual polymer filaments were capable of directly-capturing other polymer-filaments.

Using the mixing-flow chamber previously described, we examined the properties of the addition of Cpl to the MMCat/MMSc/DzII-mix (+DzII/+Cpl). Solutions of 0.8 µM [MMCat/MMSc], and 1.2 µM [DzII] mixed with 4.2 µM [Cpl] were loaded onto channels 1 and 2 of the flow chamber, respectively. Upon actuation of both channels, polymer nucleation and growth were observed. The +DzII/+Cpl configuration seeded fewer polymers compared with -DzII/-Cpl, (Figure S16), but also, remarkably, in the +DzII/+Cpl driven formation of polymer-mesh networks (**Figure 5b**, Movie 9). This resulted in the exponential growth of total polymer length per seeded filament compared with the monotonous-growth measured for -DzII/-Cpl (**Figure 5c**).

Simultaneously, the branching per filament increased faster in +DzII/+Cpl, compared to the rate observed in -DzII/-Cpl (**Figure 5d**). The simultaneous increase in total and mean filament length values (Figure S16), and in branching, suggests that the combination of DzII and Cpl promotes the emergence of a dynamic, branched polymer-mesh-network. Cleavage and subsequent incorporation of polymers into neighbouring coupled-filaments is consistent with a mechanism in which DzII-mediated cleavage and Cpl-mediated inter-filament capture jointly remodel and expand the network. Polymer fragments remain tethered in place by other Cpl-coupled polymers, exposing newly-formed filament-ends from which more polymer can grow, thus sustaining the autocatalytically-accelerating growth of the mesh-network.

## Conclusions

We have transformed a thermodynamically controlled DNA polymer into a dissipative system that polymerizes and depolymerizes spontaneously, driven by covalent-bond cleavage and monomer consumption. Held in a non-equilibrium steady-state, the filaments show emergent behaviours at two levels: dynamic instability in individual filaments and autocatalytic mesh formation in the bulk.

By containing a switchable internal-state with an intrinsic lifetime, the hybrid monomers polymerized and depolymerized spontaneously and autonomously as a result of the MMCat cleavable-RNA insertion-induced defects in the polymer-lattice.

We further expanded the properties of the system by the insertion of catalyst binding-domains within the MMCat unit, allowing for the catalytic, controlled switching (cleavage) of the internal-state within the MMCat. By using an inter-molecular source of catalytic activity, Dz(I-IV), we were able to explore a phase-space beyond the intrinsic RNA-cleavage activity of the hybrid polymers, and most importantly, pinpoint the kinetic conditions which allowed for the emergence of dynamic instability-like properties.

Due to the system properties depending tightly on MMCat/MMSc concentrations, Origami-seed-functionalized mixing-flow-chambers were used for maintaining constant MMCat/MMSc concentrations, and allowed observing the system for longer time-periods. We defined three different regimes in which the system displayed different properties: (i) When [DzII] << 0 - 0.9 µM, rapid polymerization and filament formation was observed, with a rate of 0.3 ± 0.1 µm/min for -DzII. (ii) When [DzII] = 1.2 ± 0.3 µM, polymer formation is observed upon flowing of the MMCat/MMSc/DzII-mixture. Polymerization is followed by the DzII-induced cleavage and subsequent recovery of the individual polymer filaments by assimilating active-MMcat units onto filaments. These counterweighted growth and collapse rates led to dynamic transitions between growth and collapse, with ca. 10% of filaments going through multiple cycles, resembling dynamic instability. (iii) When [DzII] >> 1.5 µM, cleavage dominated, and sparse polymerization was accompanied by rapid depolymerization and filament depletion due to the increased catalyst activity. By lowering [DzII] to 1.2 ± 0.3 µM, we were able to rescue the dynamic instability phenotype.

Since the DzII-catalyst is freely diffusing, catalyst/filament interactions can occur along the entire filament and result in scission at internal locations, and that these synthetic polymers lack the ageing feature of tubulin. Reconstituting this ripening feature would be useful in developing intramolecularly catalytically-state-switching synthetic filaments.

Attempts were also made at transforming the system into a closed system by means of ligating spent-MMCat. Although the ligation reaction was capable of ligating the spent MMCat in the bulk of a test tube, imaging of the closed system was not successful, and will be pursued further.

We also evaluated the cooperative properties of the polymer. By adding the coupling strand Cpl to the MMCat/MMSc/DzII mix, polymers were now capable of directly capturing neighbouring filaments. The Cpl induced capture led to autocatalytic growth, in which polymer mesh formation led to larger polymer-networks forming. These mesh structures displayed an exponential correlation between filament nucleation events and total polymer length, compared with the direct correlation observed without the DzII catalyst and Cpl.

Dynamically unstable, autocatalytic mesh-forming polymers could help elucidate complex cytoskeletal phenomena and serve as a flexible, bottom-up cytoskeleton for life-like systems combining the so-called generic-to-life principles: dissipation, autocatalysis (replication), steady-state formation and learning.

## Methods

### Preparation of DNA and DNA/RNA monomer units

Monomer units used were of the DAE-E type, and their sequences derived from the previously reported nRE/nSE configuration. DNA strands were ordered from IDT (Leuven-Belgium/Coralville, USA) in desalted or HPLC-purified form (see table). Chimeric DNA/RNA strands were ordered RNase-free HPLC-purified. All DNA and DNA/RNA strands were dissolved using RNase free water in a laminar hood to a concentration of 200 μM. Concentrations were then verified using a nanodrop-1000 spectrophotometer (Thermo Fisher), and final concentrations of the nucleic acid strands were adjusted accordingly to 100 μM. Equimolar amounts of the strands composing (**MMCat**: Cat1, Cat2-Sub2R-Polar, Cat3F-Overhang, Cat4, and Cat5-Subcomp, **MMSc**: Sc1, Sc2-Polar, Sc3F, Sc4 and Sc5) each monomer-type were mixed to form 20 μM [MMSc] or 16.7 μM [MMCat] monomer stock solutions. The different monomer stock solutions were then aliquoted and stored in -20 °C for up to 5 weeks, assessed for RNA cleavage prior to use using urea-PAGE, and quantified for the amount of cleaved RNA strands.

### DNA or DNA/RNA bulk filament polymerization assay

DNA polymer formation assays were performed using different monomer variants (Table S1). New premixed monomer stock solutions were thawed and diluted to 5 μM in freshly mixed, degassed, and filtered 1xTAE-Mg^2+^ buffer, pH 8.36. Newly made monomer solutions were then annealed using an Eppendorf Themo-mix fitted with a 0.5 ml tube insert and a heated lid. Annealing was performed by incubating for 5′@92 ℃, followed by cooling at a rate of 0.1 ℃/min to 21 °C overnight. Monomer formation was corroborated using 10%, 19:1 native 1xTBE-Polyacrylamide gels. Monomer-loss from the annealing phase was estimated at ca. 25% of total monomer concentration using urea-PAGE (not shown). Monomer stocks were mixed in DNA LoBind tubes (Eppendorf) and diluted to concentrations between 25 nM to 1600 nM in 1xTAE-Mg^2+^ buffer and 2 mg/ml BSA, depending on the assay performed. Stocks were newly made prior to each experiment.

### Determining of monomer critical concentration

Monomer stocks for the all-DNA or hybrid DNA/RNA systems were prepared as previously described. A gradient of the individual monomer stocks was mixed in the following concentrations: 30 nM, 39 nM, 78 nM, 156 nM, 312.2 nM, 625 nM, 1250 nM, and 2500 nM. Each two matching stock concentration were then mixed at 21 ℃, ensuring minimal polymerization activity, had their absorbance measured at 260 nm using a nano-drop nd-1000, centrifuged at 20,000 g for 5′, and their supernatant measured for absorbance at 260 nm, to evaluate the initial concentration of free monomer in solution, ***[M_0_]***. After the base-line measurements were obtained, the individual concentrations of monomer-mix solutions were let incubate at 35 °C for 90’ in an Eppendorf Themo-mix fitted with a 0.5 ml tube insert and a heated lid, to avoid condensation formation on the lids of the tubes. After the 90’ incubation period, the monomer mix tubes were centrifuged at 20,000 g for 5′, in order to precipitate the formed polymer structures. At higher monomer concentration, a visible pellet was seen at the bottom of the tubes. The supernatant was measured for absorbance at 260 nm, and the remaining free monomer in solution was evaluated, ***[M_t_]***. The polymerized monomer fraction, ***[M_p_]***, was calculated using equation 1.

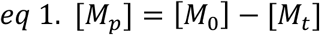

Samples were then verified for presence or absence of polymer formation by loading 1 μl of monomer-mix onto a 2mg/ml BSA coated, glass slip/slide sandwich, and observing the fluorescently (Alexa488) labelled polymers.

### Measurement of Michaelis-Menten kinetic parameters

A 10 µM stock solution containing the MMCat and MMSc in 1xTAE-Mg^2+^ buffer was annealed overnight as previously described. A dilution ladder consisting of 7 concentrations of the annealed polymer solution was made, consisting of the following concentrations: 5 µM, 2.5 µM, 1 µM, 0.5 µM, 0.2 µM, 0.1 µM, and 0.05 µM. Keeping the polymer solution at 21 ℃, to ensure negligent catalytic activity, the DNAzyme variant was then pipetted into the polymer stock, resulting in a final 10 µM [DNAzyme] and a volume of 20 µl. 2 µl of the polymer/Dz mixture was then mixed with 8 µl of ice-cold stop buffer (80% w/v deionized formamide, 10 mM EDTA (pH 8.0), and 1xTBE) and designated as t = 0. The remaining sample was then quickly loaded onto an Eppendorf Themo-mix set at 35 ℃, and 2 µl aliquots were removed and deactivated at the desired time points, up to t = 30’. The time point aliquots were then mixed with 10 µl of 2x Gel tracking buffer to yield a 10x dilution, and diluted again 10-fold using the stop buffer. 18 µl of the 100-fold diluted time points were then loaded onto a pre-run 1mm thick 15 %, 19:1 cross-linked ,7M urea, 1xTBE, PA gel, and let run for 90’ at 200V. The resulting gels were then scanned at 488nm and 640 nm using a Bio-rad bio-imager, and observing the fluorescently-labelled substrate and product bands. Gels were normalized between themselves using an equimolar substrate-well pipetted to all gels. The initial-rate region of the substrate formation bands was then plotted for all the different substrate (polymer) concentrations, and fit using Matlab.

### Insertion of modifications to DNA and DNA/RNA monomer strands

To avoid the formation of steric hindrances along the filament, all modifications inserted to the different nucleic acid strands comprising the different monomer variants were added on the outside facing ends of the nucleic acid stands, in relation to the polymer structure. i.e. the modifiable terminals were the 3′ ends for strands Sc1, Cat1, Sc5, and Cat5, the 5′ ends for strands Sc2, Cat2, 4Sc, and Cat4, and both the 5′ and 3′ ends for strands Sc3 and Cat3. The incorporation of the RNA bases on the Cat2 strand was done using similar considerations, which made the DNAzyme recognition site accessible to the diffusing DNAzyme-units in the bulk outside the polymer.

### Glass slides and slips functionalization and passivation for DNA or DNA/RNA polymer imaging

Imaging experiments were performed on passivated, and biotinylated glass. To better control the interaction of the filament and the filament monomers with the glass surface, all glass surfaces were cleaned and passivated. A 3M NaOH solution filled slip or slide holder was loaded with untreated glass and left to etch for 60’. Glass was then removed from the NaOH solution and rinsed three times using Millipore ultrapure-water, and dried using N_2_. Cleaned glass was then plasma-cleaned (Plasma Technology GmbH, Herrenberg-Gueltstein, Germany) for 3′ in a partial atmosphere of 0.35 mbar O_2_, and pipetted with 25 μl of freshly-made 1 mg/ml Silane-PEG_3500_-Biotin (CD Bioparticles, USA) dissolved in dry DMSO. Solution-wetted glass was then sandwiched with a matching glass piece, and let incubate for 45’ at 70 ℃. Glass sandwiches were subsequently separated, washed three times with Millipore ultrapure water, and dried using N_2_. Prior to imaging, the biotinylated glass was passivated using a mixture of 1% F127 Pluronic detergent, 0.1 mg/ml blocker casein (Thermo Fisher) in 1xTAE-Mg^2+^ and left to incubate for ca. 45’. Glass was rinsed using Millipore ultrapure water, and dried using N_2_. For experiments involving the binding of nucleic acids to the glass surface, a solution of 0.75 μM Neutravidin was prepared in 1xTAE-Mg^2+^ and let to incubate on the objective-facing glass piece for 5′, followed by washing of unbound neutravidin using 1xTAE-Mg^2+^.

### Flow chamber assembly and preparation for TIRF imaging

A two-part, PDMS-resin was mixed according to kit instructions (SYLGARD 184, Dow corporation, USA), degassed for 75’, and poured onto a silicon wafer imprinted with the flow chamber design. The PDMS-resin was let to cure overnight at 70 ℃. The following day, the fully cured PDMS-chip was cut to individual flow chamber chips out of the PDMS mould using a scalpel. Inlet and outlet holes were punched into the PDMS chips using a 0.75 mm biopsy punch. Flow chamber chips were then stored for up to 2-months in a closed, desiccated, glass container. Prior to assembly, the flow chamber chip surface was lightly cleaned using scotch-tape. A 24×60 mm H1.5 glass slip (Marienfeld, Germany) was lightly etched using a 3M NaOH solution bath for 60’, followed by rinsing three times with Millipore ultrapure water and dried using N_2_. Cleaned glass was then plasma cleaned for 3’ in a partial atmosphere of 0.35 mbar O_2_, and pipetted with 50 μl of 1 mg/ml Silane-PEG_3500_-Biotin (CD Bioparticles, USA) dissolved in dry DMSO. Solution-wetted glass was then sandwiched with a matching glass piece, and let incubate for 45’ at 70 ℃. Following the biotinylation of the glass surface, the intended imaging field of view in the glass was marked (on the outside) with a permanent marker and masked using a flat, cured PDMS block. Matching PDMS chip and glass slip pairs were then plasma cleaned for 3’ in a partial atmosphere of 0.35 mbar O_2_. The PDMS blocker was then removed from the glass, and the glass slip and PDMS chip were sandwiched together quickly to avoid water adsorption to the hydroxylated surface, using the previously drawn marks to align the flow chamber channel with the glass slip. The entire PDMS-glass device was then let to incubate at 45 °C for 75’. Flow chamber assemblies were then stored in a closed container. Prior to imaging, the flow chamber was assembled by plugging 0.6 OD mm PTFE tubing (Bohlender, Germany) to the used inlets and outlet. Syringes tipped with 30G blunt-needles, and infused with 1ml 1xTAE-Mg^2+^ buffer solution were then loaded onto Harvard Apparatus 11 Elite syringe pumps. The pumps were set to a flow rate of 10 μl/min and let run for 10’. Following buffer rinsing, one of the pump channels was refitted with a syringe containing 0.3 ml of 1% F127 Pluronic detergent, 0.1 mg/ml blocker casein (Thermo Fisher) in 1xTAE-Mg^2+^, and let run for 10’ at a rate of 10 μl/min. After passivation, the channel was washed with 1xTAE-Mg^2+^ for 10’ at a rate of 10 μl/min. Following the wash, a syringe containing 0.3 ml of 0.75 μM Neutravidin in 1xTAE-Mg^2+^ was loaded onto the pump and let run for 10’ at a flow rate 10 μl/min, and washed afterwards using 1xTAE-Mg^2+^ for 10’ at 10 μl/min. Following the neutravidin coating step, a syringe containing 0.3 ml of 1:5000 dilution of DNA-Origami seed stock in 1xTAE-Mg^2+^ was loaded onto the pump and let run for 2’ at a flow rate 10 μl/min, and washed afterwards using 1xTAE-Mg^2+^ for 10’ at 10 μl/min. Pump channel(s) were then loaded with appropriate sample containing syringes.

### DNA or DNA/RNA filament polymerization imaging using epifluorescence microscopy

Newly diluted 5 μM Monomer stocks were mixed in DNA LoBind tubes (Eppendorf) and diluted to 25 nM, 100 nM, 200 nM, or 400nM in 1xTAE-Mg^2+^ buffer and 2 mg/ml BSA and let incubate for 15’’ at 35 ℃. 2 μl of monomer mix was loaded onto cleaned, untreated 1.5H (Marienfeld, Germany) 22×22 mm kept at 35 °C and sandwiched onto a glass slide heated to 35 ℃. The sample sandwich was loaded onto an atmosphere and temperature-controlled Olympus IX-81 microscope fitted with a x60 1.4 NA oil-immersion objective and a Hamamatsu ORCA Quest sCMOS camera sensor and a Lumencor Spectra X LED module. Timelapses of filament polymerization were captured using 1’’ intervals and 100 msec exposure time at the excitation wavelength of 488 nm, or 647 nm. Timelapses were then processed using FIJI, using custom macros.

### DNA or DNA/RNA polymerization imaging using TIRF microscopy

Newly diluted 5 μM Monomer stocks were mixed in DNA LoBind tubes (Eppendorf, Germany) and diluted to 400 nM, 800 nM, or 1.2 μM in 1xTAE-Mg^2+^ buffer and 2 mg/ml BSA and let incubate for 15’’ at 35 ℃. The custom built, Olympus IX81-based, TIRF microscope was equipped with a Hamamatsu Orca Fire sCMOS camera, four diode-laser sources: 405 nm, 561 nm, (Coherent, Inc. Ca, USA), 488 nm, and 647 nm (Toptica Photonics, Germany), a 60x 1.4NA, temperature-corrected, oil-immersion objective heated with a Bioptechs objective heater, a Pecon Tempcontrol 37-2 controller and heated stage, and two Harvard Apparatus 11 Elite syringe pumps, allowing for the individual control over the two solution channels. Temperature was maintained at 35 °C for the duration of the imaging assays. The microscope was operated using a custom-made micro-manager interface. Time lapse acquisition rates were set between 25 milliseconds to 1 seconds, depending on the experiment being run, for a duration of 5’ to 240’. Slip/slide sandwich experiments were performed as previously described, with the modification of including a 7 μm thick double-sided tape (PolyK Technologies LLC, USA) between the slip and the slide, to allow for higher sample volume and longer imaging time.

### Flow chamber imaging assays using TIRF microscopy

A 400 μl solution of 2 mg/ml BSA, and 400 nM or 800 nM MMCat/MMSc Monomer-mix was loaded onto a 30G blunt-needle tipped syringe and placed in syringe pump 1. Flow rate was kept at a total of 5 ± 1 μl/min between the two channels, with the ratio between the channels depending on the experimental conditions required. The TIRF microscope was used as previously described. Flow chambers were assembled and functionalized as previously described. The imaging field of view was chosen within the DNA-Origami seed coated region of the flow chamber. Upon loading of the monomer-mix solution filled syringe into pump-channel 1, flow was commenced at a rate of 5 μl/min. Upon reaching the Origami-seed-modified imaging area, monomer conjugation onto the seeds and polymer formation was observed. The flow rate was kept at 5 ± 1 μl/min for the duration of the imaging assay.

To observe the dynamic instability property, a 30G blunt-needle tipped syringe loaded with a solution of 400 μl of 2 mg/ml BSA, and 2.4-4.8 μM DzII catalyst was placed in pump-channel 2 and set at 4.5 μl/min. The flow chambers utilized a four-inlet design, with channels 1 and 2 meeting at a mixer, to produce a homogenous distribution of components within the flow chamber. Subsequently, pump-channel 1 was set at 0.5 μl/min, in order to avoid bubble formation within the flow chamber. After ca. 120’’, the pump-channel 1 was set at 3 ± 0.5 μl/min and channel 2 was set a flow rate of 2 ± 0.5 μl/min. The evolution of the individual polymers over time was then recorded at 488 nm using the Alexa488-labelled MMSc monomer, and at 640 nm using the Atto 647 -abelled MMCat. Experiments utilizing the DNAzyme-catalyst DzII and the Cpl coupling strand were performed using channels 1 (MMCat/MMSc monomer-mix) and the direct-injection inlet 3 (Cpl or DzII/Cpl mix). In a 30G blunt-needle tipped syringe, 400 μl of a mixture of 2 mg/ml BSA, containing either 4.2-8.4 μM Cpl Coupling strand or both 2.4 μM DzII catalyst and 8.4 μM Cpl Coupling strand mixed together, and the syringe was then loaded to pump-channel 3. Subsequently, pump-channel 1 was set at 0.5 μl/min, in order to avoid bubble formation within the flow chamber. After ca. 120’’, the pump-channel 1 was set at 2.5 ± 0.5 μl/min and channel 3 was set a flow rate of 2.5 ± 0.5 μl/min. The evolution of the polymers and the mesh formed over time were then recorded at 488 nm using the Alexa488 labelled MMSc monomer, and at 640 nm using the Atto647 labelled MMCat.

